# Analysis of arbuscular mycorrhizal fungi (AMF) community composition associated with alkaline saline sodic soils

**DOI:** 10.1101/2024.02.01.578375

**Authors:** N Marquez, JM Irazoqui, MB Ciacci, AF Amadio, FD Fernandez, ML Giachero

## Abstract

Marginal soils affected by salinity, sodicity and alkalinity decrease crop productivity. In this context, a viable alternative strategy lies in the remediation of degraded lands using beneficial microorganisms. This study aims to characterize native arbuscular mycorrhizal fungal (AMF) communities by sequencing PCR amplicons that cover most of the small subunit rRNA (SSU) gene, the complete internal transcribed spacer (ITS) region, and a portion of the large subunit (LSU) rRNA, employing Oxford Nanopore Technologies (ONT). Three field sites, with varying crop conditions, were selected: a patch with no crop growth (Site 1), a patch with corn stubble (Site 2), and a patch with wheat plants exhibiting 15 days of growth (Site 3). Soil analyses revealed distinct characteristics - alkaline saline sodic soil (ASS) for Site 1, moderately alkaline soil (A) for Site 2, and neutral soil (N) for Site 3. ONT sequencing yielded a total of 4,040,470 raw reads from which 19.13% survived after quality and length filter. Reads were grouped in 556 clusters, of which 222 remained after bioinformatic analysis. Despite moderate error rates in 9.4.1, flowcells chemistry, using a clustering and polishing approach facilitated the ecological analysis and allowed a better taxonomic resolution. Bioinformatic analysis showed no significant differences in AMF diversity among soils. However, results suggest the dominance of *Glomeraceae* and *Acaulosporaceae* families, specifically the genera *Glomus* and *Acaulospora* in ASS soil. Further exploration is required to better understand their role in promoting plant growth under adverse conditions. The study highlights the significance of cutting-edge sequencing tools in advancing the comprehension of essential symbiotic relationships for sustainable agriculture in challenging environments.

## INTRODUCTION

The accumulation of water-soluble salts is one of the forms of soil degradation that impact over 15% of the total land area of the world. According to the GSASmap (Global Map of Salt-Affected Soils - FAO), based on the current information from 85% of global land area, more than 424 million hectares of topsoil (0-30 cm) and 833 million hectares of subsoil (30-100 cm) are salt-affected ([GSASmap](https://www.fao.org/soils-portal/soil-survey/mapas-historicos-de-suelos-y-bases-de-datos/global-map-of-salt-affected-soils/es/)). Moreover, the soil’s pH constitutes the primary indicator of nutrient availability for plants, exerting significant influence on the solubility, mobility, and accessibility of various constituents within the soil matrix. The pH of soil ranges from 3.5 (highly acidic) to 9.5 (highly alkaline), with a pH of 6.5 considered ideal ([GSASmap](https://www.fao.org/soils-portal/soil-survey/clasificacion-de-suelos/sistemas-numericos/propiedades-quimicas/es/)). Often, alkaline soils occur simultaneously with salinity conditions, resulting in the formation of another category of soils (commonly referred to as sodium saline soils); additionally, in some cases, saline soils can transform into alkaline soils, and vice versa (Naidu and Rengasamy, 1993). However, because leaching may have occurred previously, alkali soils do not always contain excess soluble salt. Saline and alkali soils are conceptually classified according to the electrical conductivity (EC), exchangeable sodium percentage (ESP), and pH. Sodium saline soils have an EC greater than 4 dS m-1 (∼40 mM NaCl) at 25°C, ESP greater than 15%, and a pH between 8.5 and 10 (Rengasamy, 2010).

The effect of salinity stress on crop physiology and metabolism is complex, but generally reduces available water, decreasing plant growth by an osmotic effect, and produces nutritional disorders and toxicities in the plant cells (Munns 2002; Kundu 2022). These physiological stresses can affect important processes, such as photosynthesis, lipid metabolism and growth among others, thereby negatively affecting crop productivity. Marginal soils due to salinity and sodicity have detrimental effects not only on agriculture but also on the microbial populations in soil. In fact, microbial community structure and activities are potential indicators to assess the severity of the land degradation (Singh 2016). One of the primary goals of agriculture is to meet the food demand while minimizing environmental impact, and a promising approach to achieve this is by expanding agricultural soils to marginal and degraded lands. In this context, a viable alternative strategy for increasing crop production while mitigating the adverse effects on the environment lies in the remediation of saline and sodic soils using beneficial microorganisms. The soil microbiome could assist plants in overcoming abiotic stress, thereby promoting sustainable agricultural practices (Singh et al., 2011; Qin et al., 2016; Chaudhary et al., 2023). Although soil microorganisms play a central role in nutrient cycling, there is still a lack of comprehensive understanding about the responses and adaptations/remodeling of microbial communities under varying salinity conditions (Zheng et al., 2017).

Mycorrhizae are symbiotic associations between certain soil fungi, the Arbuscular Mycorrhizal Fungi (AMF), and the roots of most terrestrial plants (Gianinazzi et al., 2010). These beneficial symbiotic associations may have functional diversity potentially implicated for agroecosystem management (Verbruggen & Kiers, 2010). It is well known that under stress conditions, the major role of AMFs is to alleviate the limitation in plant growth. The implicated mechanisms are, for example, the improvement of molecular and physiological processes in the host plant (photosynthesis, efficient use of water, production of osmoregulators) and/or the enhancement of plant nutrient acquisition (Porcel et al., 2012; Hashem et al., 2019; Hsieh et al.,2022). However, mycorrhizal communities can also be affected by environmental factors and host plant type (Aliasgharzadeh et al, 2001). Salinity could delay the germination of spores in the soil and directly affect hyphal growth, resulting in a reduction of mycorrhization in their host plants (Jahromi et al., 2008; Delvian and Rambey 2019). Thus, considering that the community structure of AMFs will depend on chemical and physical properties of soil, it’s important to know the diversity and abundance of these fungi under multiple field conditions and their role in stress alleviation (Kumar et al., 2015). Several results indicated that indigenous AMF communities adapted to certain edaphic factors may have advantages compared with non-native AMF (Lambert et al., 1980; Rua et al., 2016). For instance, *Glomus geosporum*, an arbuscular mycorrhizal fungus (AMF) isolated in Europe from saline, sodic, and gypsum-rich soils, exhibited remarkable adaptability to these challenging conditions and demonstrated significant potential in enhancing plant resistance against salinity (Landwehr et al., 2002). Over the course of several years, this AMF showcased the capability to significantly enhance the growth of various cucurbitaceous plants, pea, and rice under salt stress conditions (Okon et al., 2019; Meça et al., 2016; Parvin et al., 2019; Tisarum et al., 2020). Likewise, recent studies stated that grass plants (*Lasiurus scindicus*) inoculated with AMF isolates from saline soils in Saudi Arabia exhibited better growth parameters compared to plants inoculated with strains from non-saline soils (Malik et al., 2022).

The identification of soil microbial composition associated with marginal soils is crucial for comprehending their ecosystemic role and can potentially facilitate soil remediation or enhance host plants adaptability to adverse conditions. Hence, the primary aim of this study is to characterize native arbuscular mycorrhizal fungal communities in alkaline, saline, and sodic soils by Oxford Nanopore Technologies (ONT) sequencing of Glomeromycota rDNA. In this way, sequencing will provide affordable AMF identification and will allow analysis of the different indigenous taxa adapted to adverse conditions potentially implicated in crop protection from a biotechnological perspective.

## MATERIAL AND METHODS

### General conditions of the Field sites

The study was conducted in Punta de Agua, a location in Córdoba province, Argentina. In the research field, conservationist management across all study plots was implemented, incorporating rotations, cultivation of service crops, and the application of nanofertilizers. The region exhibits a temperate thermal regime (mesothermal) with an annual average temperature of 17.3 ºC. Notably, January records the highest temperature at 23.8 ºC, while July experiences the lowest at 11 ºC, resulting in an annual thermal amplitude of 12.8 ºC (INTA, 2008). The first frosts typically occur around May 15, with the last frosts observed by September 15. Frosts may deviate by 15 to 20 days from the average dates. A significant feature is the frost-free period lasting approximately 240 days (Capitanelli, 1979).

Soil composition is predominantly Natraqualf, categorized based on Soil Taxonomy (Soil Survey Staff. 1999) and exhibits variations primarily in texture and drainage across different cartographic units (Jarsún et al., 2003). In the specific location of Punta de Agua, we find 12,000 hectares with moderate salinity and an additional 11,000 hectares characterized by superficial alkalinity. These soils suffer from imperfect natural drainage, leading to frequent flooding. Notably, the surface horizon of these soils is discolored, and they possess a high exchange capacity (Georgas et al., 2003).

### Field sites and soil samples selection

Three patches with irregular crop growth and different properties were selected as study sites within the field: at site 1 a patch of no crop growth was observed (32º33.5’32. “S 63º43.9’54 “W - Figure 1A); site 2 featured corn stubble (32º33.5’49 “S 63º43.9’56 “W - Figure 1B), and site 3 featured a patch with wheat plants exhibiting 15 days of growth (32º34’42. “S 63º44.1’41 - Figure 1C). Soil samples were collected from these three visually different patches at a depth of 0-30 cm using shovels. The composite soil sample from each site consisted of a pool of three randomly chosen points, separately 300 cm from each other, which were combined and mixed. Samples were transferred within black plastic bags and maintained at 25ºC for physical and chemical analyses. For the AMF community analysis, each composite soil sample was homogenized, sieved to remove plant debris, and stored in 50 ml plastic tubes at room temperature.

**Figure 1:**
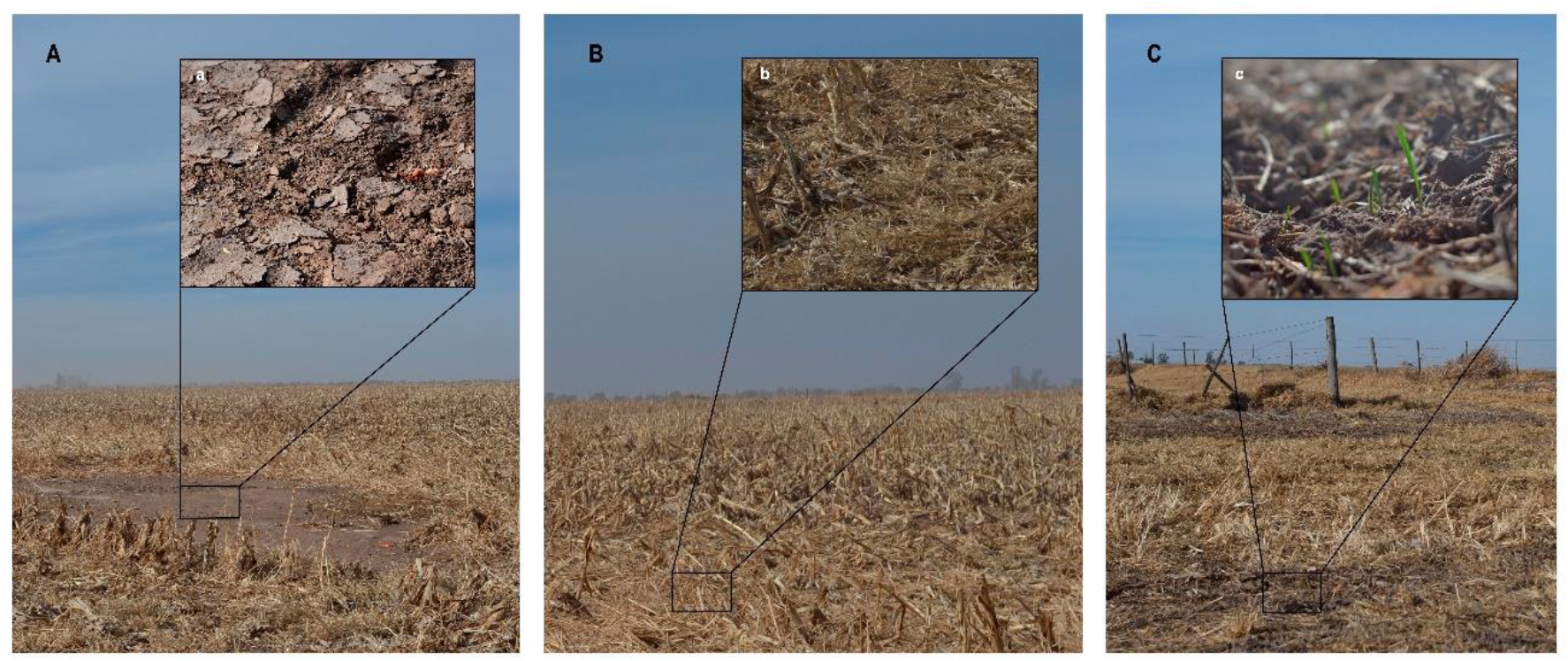
Sampling Sites. (A) site 1: a patch of no crop growth; (B) site 2: featured corn stubble; (C) site 3: wheat plants exhibiting 15 days of growth. In Figures a, b and c the sampled soil area for each point is detailed.

### Soil physicochemical properties

The physicochemical properties of each soil type were measured. Parameters like pH, electrical conductivity (EC), total nitrogen (Total N) were determined by the semi-micro Kjeldahl method (SAMLA, 2004), organic carbon (OC) by oxidation with potassium dichromate solution, available phosphorus (P) by turbidimetry, potassium (K), calcium (Ca), magnesium (Mg) and sodium (Na) extracted with ammonium acetate solution, and cation exchange capacity (CEC) according to Van Raij (1998). For each soil type, two subsamples were taken for DNA extraction.

### DNA extraction, PCR amplification and sequencing

Total DNA was extracted from 0.25 g of soil using the DNA PowerSoil PRO KIT (QIAGEN) according to the manufacturer’s instructions. For each soil sample, two different extractions were done: neutral soil (N1+N2), alkaline soil (A1+A2) and alkaline saline sodic soil (ASS1+ASS2). Concentration and purity of total DNA were determined in a ND 1000 UV–vis spectrophotometer (NanoDrop Technologies).

DNA extracts were diluted 10-fold and used as templates to amplify the target fragment of Glomeromycota rDNA to compare and describe AMF communities between soil samples (Kolaříková et al., 2021). AMF-specific primers (Supplementary Table 1) developed by Simon et al. (1992), Lee et al. (2008), Kruger et al. (2009) and Kolaříková et al. (2021) were used.

Two PCRs were performed using the following primer combinations: NS31/LSUmAr or AML1/LSUmAr. PCR reaction mixture (25 µl) included Q5 reaction buffer (5X), 0.4 mM of each dNTP, 0.5 µM of each primer, 0.02 U/µl of the Q5 High Fidelity DNA polymerase (New England BioLabs) and 1 µl of the template. The amplification cycle was done in an Eppendorf Mastercycler Gradient (Eppendorf, Hamburg, Germany) with the following parameters: 98°C for 5 min, followed by 40 cycles at 98°C for 10 s; at 65°C for 30 s; at 72°C for 90 s, and a final extension at 72°C for 5 min. A negative PCR control, utilizing water as the template, was incorporated into the sample set.

PCR products from these initial PCRs using primer combinations (NS31/LSUmAr or AML1/LSUmAr) were pooled, diluted 250-fold, and used as template for a second PCR using the following primers: NS31_Glo3/LSUmBr. PCR reaction mixture and cycling conditions were the same as in the initial PCRs except that the annealing temperature was 68°C and 30 cycles were done. This second PCR was performed in replicates.

The resulting amplicon (2.5 Kb long) of the nested PCR, included a large part of the small subunit rRNA gene (SSU), the complete internal transcribed spacer region (ITS1–5.8S–ITS2) and part of the large subunit rRNA gene (LSU).

The replicates of the second PCR were pooled for each sample and purified with Agencourt AMPure XP beads (Beckman Coulter) according to the manufacturer’s instructions. Amplicon concentration was quantified with a ND 1000 UV–vis spectrophotometer (NanoDrop Technologies) and Quantus (Promega, USA).

Sequencing libraries were prepared using the Ligation Sequencing Kit (SQK-LSK109) and the Native Barcoding Expansion 1-12 (EXP-NBD104) following the manufacturer’s instructions provided by Oxford Nanopore Technologies. In brief, the pooled amplicons were initially repaired using the Ultra II End Repair / dA-tailing Module (NEB, USA). Subsequently, barcodes were ligated using the NEB Blunt/TA Ligase Master Mix (NEB, USA). The entire sequencing procedure was performed on a MinION Mk1B device with a flowcell R9.4.1. All sequencing data were deposited at NCBI Genbank under Bioproject accession PRJNA1062293.

### Bioinformatic analysis

All FAST5 files were basecalled using Guppy (v6.3.2) with the super accurate model. Resulting reads were filtered by length (minimum length allowed 2000 bp, maximum length allowed 3200 bp) and by quality (read average quality >= 12). All remaining reads from all samples were combined and clustered using isONclust (Sahlin & Medvedev 2020) with options --min_fraction

0.9 –mapped_threshold 0.9. For each cluster, a consensus sequence was obtained using abPOA (Gao et al., 2021) and then polished using medaka (https://github.com/nanoporetech/medaka). Clusters with only one read or consensus sequences over 3200bp were discarded. To check if the cluster reconstruction was correct, we compared them with the primers used for the amplification using BLAST+ (Camacho et al., 2009). All reads that had one or both missing primers, or had more than 1 mismatch were discarded. Finally, the abundance table was constructed based on the number of reads in each sample that were clustered together.

With the final set of sequences, alpha and beta diversity indexes were calculated using R packages vegan (Dixon 2003) and phyloseq (McMurdie & Holmes 2013). Also, a taxonomic profile was obtained using emu (Curry et al., 2022), comparing against the Maarjam database (Öpik et al., 2010). Only hits with score over 0.95 were considered.

## RESULTS

### Soil physico-chemical properties

The physico-chemical parameters of each soil type are listed in Table 1. Site 1 was a strongly alkaline soil (pH: 8.5), saline (EC: 5.15) - sodic (ESP: 26.10%), site 2 was moderately alkaline soil (pH: 8.04), not saline (EC: 2.8) and non-sodic (ESP: 7.12%) and site 3 was neutral soil (pH: 7.01), non-saline (EC: 0.68) non-sodic (ESP: 0.51%). Hereafter site 1 will be called ASS (Alkaline Saline Sodic), site 2 will be called A (Alkaline) and site 3 will be called N (Neutral).

**Table 1:** Physicochemical parameters of each sample soil. ASS: Alkaline Saline Sodic, A: Alkaline, and N: Neutral.

|  |  | Soil samples |  |  |
| --- | --- | --- | --- | --- |
| Parameters |  | ASS | A | N |
| CE (dS m<sup>-1</sup>) |  | 5,15 | 2,8 | 0,68 |
| pH |  | 8,5 | 8,04 | 7,01 |
| CEC<br>(meq/100<br>g) | Ca<sup>2+</sup> | 10,23 | 11 | 10 |
|  | Mg<sup>2+</sup> | 4,6 | 3,49 | 4,5 |
|  | Na<sup>+</sup> | 5,8 | 1,24 | 0,09 |
|  | K<sup>+</sup> | 1,55 | 1,69 | 2,8 |
| ESP (%) |  | 26,1 | 7,12 | 0,51 |
| CEC (meq/100g) |  | 22,18 | 17,42 | 17,39 |
| Total N (%) |  | 0,14 | 0,11 | 0,12 |
| P (ppm) |  | 6,98 | 4,62 | 10,5 |
| OM (%) |  | 2,88 | 3,51 | 3,77 |
| OC (%) |  | 1,67 | 2,04 | 2,18 |
| soil texture |  | clay loam | clay loam | silty clay |

Abbreviations: electrical conductivity (EC), cation exchange capacity (CEC), potassium (K), calcium (Ca), magnesium (Mg) and sodium (Na), exchange sodium percent (ESP), total nitrogen (Total N), availability phosphorus (P), organic matter (OM), organic carbon (OC), cation exchange capacity (CEC).

### Sequencing results and community diversity

A total of 4,040,470 raw reads were obtained from which 780,226 (19.13%) survived after quality and length filter (Table 2). Reads were grouped in 556 clusters, but 202 were discarded because they were singletons. Further 132 clusters were discarded because the sequence was too long or lacked one or both PCR primers. Finally, a total of 222 clusters remained, of which 156 were present in all the soil conditions, 14 clusters were unique in neutral soils (N1 and/or N2), 15 were unique in alkaline soils (A1 and/or A2) and 4 were unique in alkaline saline sodic soils (ASS1 and/or ASS2) (Figure 2 A). Figure 2 B indicates the number of clusters shared between samples. An important number of clusters were present in all samples (100 from 222 total clusters), while 156 were present in at least one subsample of each soil type. From the 15 clusters present solely in alkaline soils, 4 were shared by both subsamples. Meanwhile, the 4 clusters unique to alkaline saline sodic soils were present in only one of the subsamples (ASS2).

**Table 2:** Sequencing and alpha diversity results. Number of reads before and after quality control, number of clusters observed and Shannon index for each sample.

| Sample | Condition | Reads | Post-filter | % survived | Clusters | Shannon |
| --- | --- | --- | --- | --- | --- | --- |
| ASS1 | Alkaline Saline Sodic | 488024 | 150190 | 30,77 | 122 | 2,63 |
| ASS2 | Alkaline Saline Sodic | 993870 | 202062 | 20,33 | 168 | 2,49 |
| A1 | Alkaline | 481342 | 97549 | 20,26 | 176 | 3,32 |
| A2 | Alkaline | 1049648 | 86980 | 8,28 | 180 | 3,78 |
| N1 | Neutral | 466980 | 131402 | 28,13 | 169 | 2,34 |
| N2 | Neutral | 560606 | 112043 | 19,86 | 187 | 3,36 |

**Figure 2:**
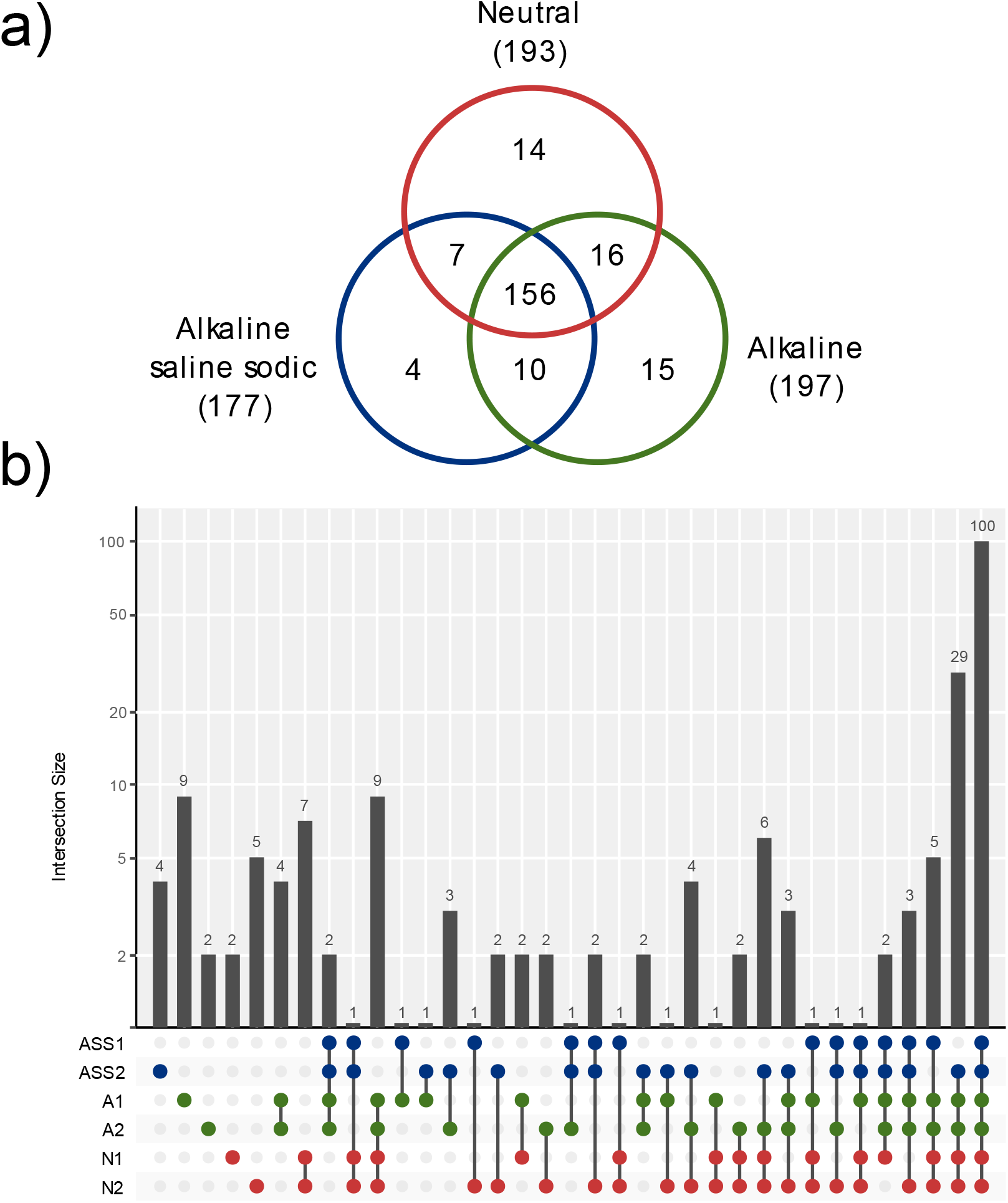
(A) **Venn diagram** showing the distribution of total cluster (n=222) between Neutral (N1 and/or N2), Alkaline (A1 and/or A2) and Alkaline Saline Sodic (ASS1 and/or ASS2) soils. (B) **Cluster distribution between samples**. The bars indicate the number of clusters shared by the different set of samples, indicated by colored dots.

Rarefaction curves showed that the sequencing effort was enough to capture most diversity in all samples (Figure 3). All samples showed a comparable number of clusters, except ASS1 that showed a lower amount, however, Kruskal-Wallis test showed that there were no significant differences between soil samples on neither richness nor Shannon diversity index (p-value > 0.05).

**Figure 3:**
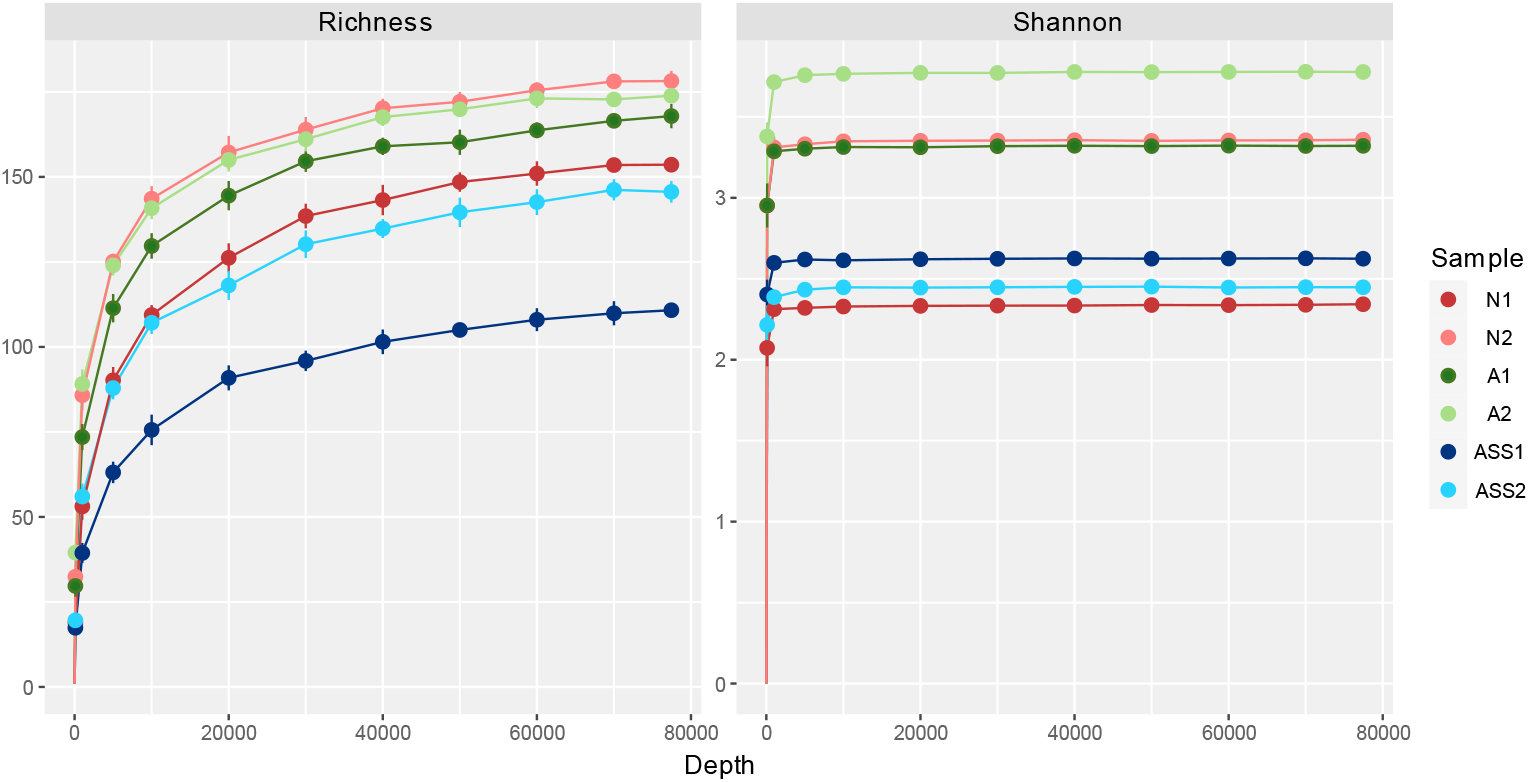
Rarefaction curves. Using both observed clusters and Shannon index, all samples reached a plateau, indicating that the sequencing effort was enough to capture most diversity in all cases.

As for beta diversity, the weighted UniFrac index showed that samples from neutral soil grouped together, as did the samples from alkaline soil (Figure 4). The same pattern was not observed in samples from alkaline saline sodic soil that were placed far apart in the DCA plot. However, no significant differences in either index were observed between soils (p-value > 0.5).

**Figure 4:**
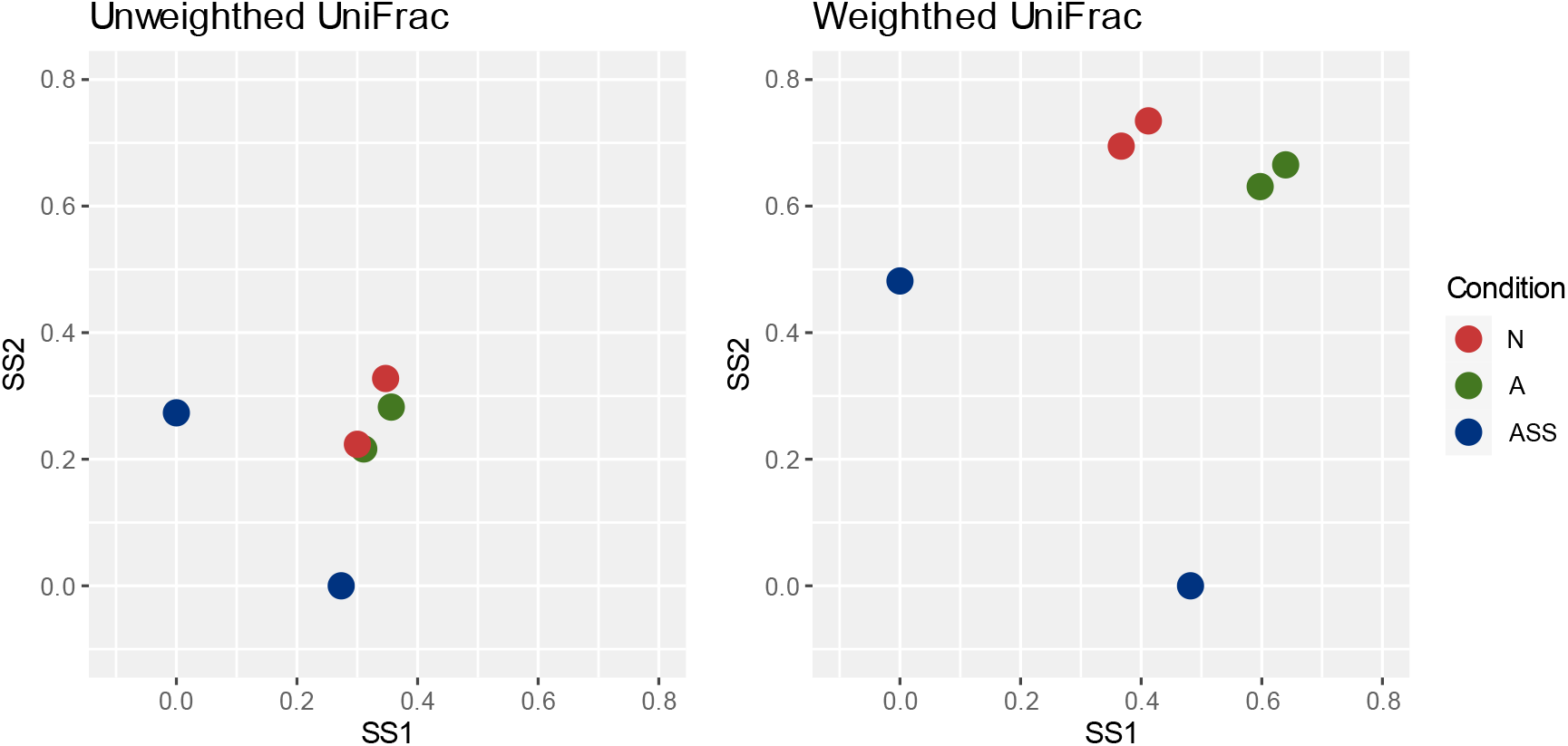
DCA using unweighted and weighted UniFrac. Beta diversity analysis using both UniFrac indexes. Using weighted UniFrac, neutral soil samples tended to group together, as did the alkaline soil samples, but the PERMANOVA analysis showed no significant differences between all soil samples (p-value > 0.05).

In all samples, the most abundant family was *Glomeraceae*, ranging between 43 and 95%. In alkaline saline sodic soils, *Acaulosporaceae* is a predominant family, adding up to 46% of ASS2, but Kruskal Wallis test showed that there were no significant differences with other soils (p-value > 0.05). Similarly, the family *Gigasporaceae* is the second most abundant family in alkaline soils, but there were no significant differences in the abundance compared to the other soil samples (Figure 5 A).

**Figure 5:**
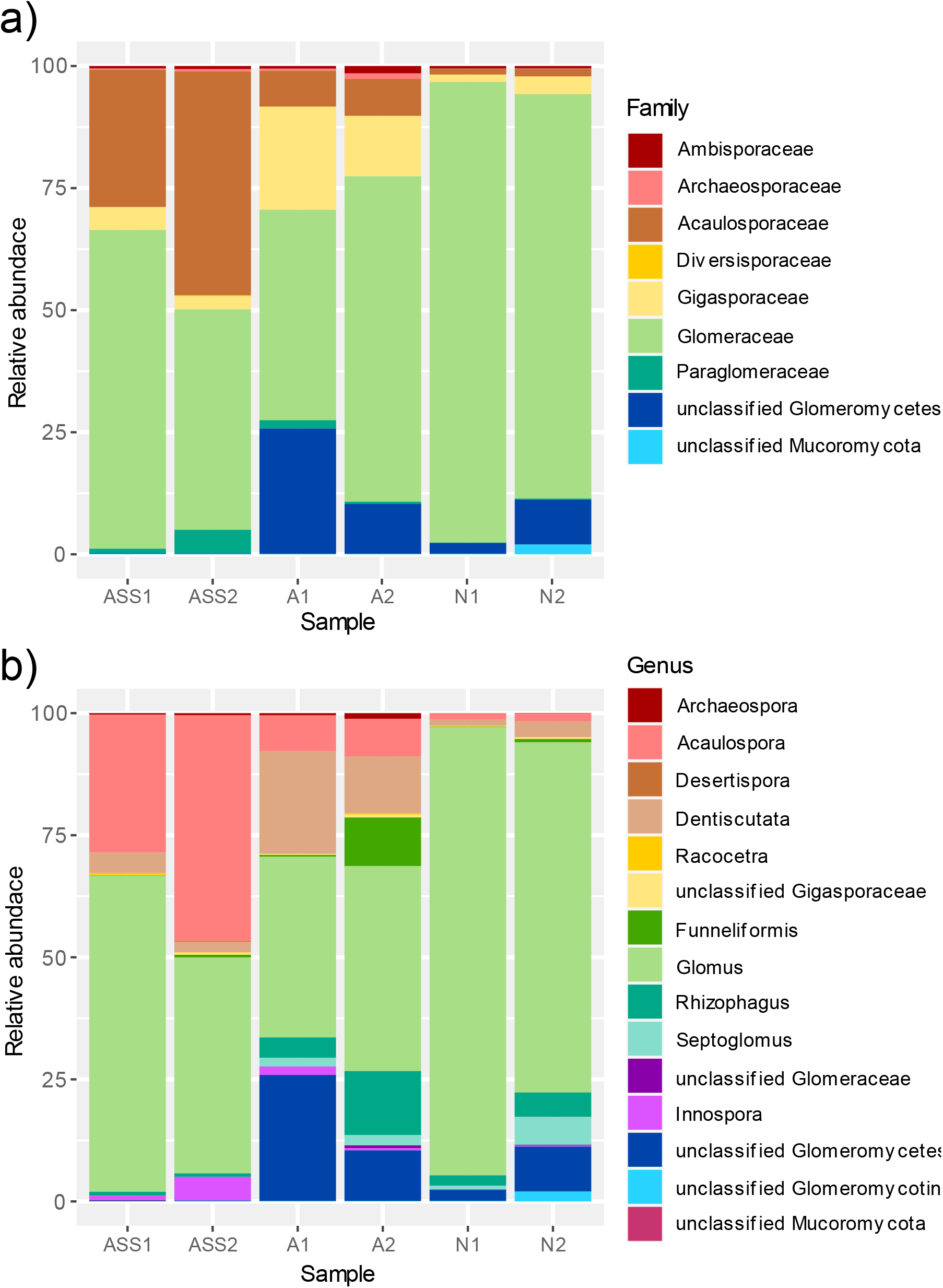
Taxonomic profiles of soil mycorrhiza. Taxonomic classification of all clusters, up to family level (A) and genus level (B), obtained by comparing with the Maarjam database.

The most representative genus in all the soils samples were: *Glomus, Rhizophagus, Septoglomus* and *Funneliformis*. In relation to the neutral soil sample, the number of clusters of the genus *Destiscutata* and unclassified members of the *Glomeromycetes* class increases in alkaline soils. In fact, the unique clusters in A soil corresponded to members of the genus *Destiscutata, Glomus, Acaulospora, Archeospora* and *Paraglomus*. Meanwhile, alkaline saline sodic soils exhibited the highest amount of *Acaulospora* and the loss of members of the genus Septoglomus was observed (Figure 5 B). The unique clusters in the ASS2 subsample were identified as *Gigaspora, Glomus* and unclassified *Mucoromycota*.

## DISCUSSION

The current study explores the community structure of arbuscular mycorrhizal fungi (AMF) in three selected sites where imperfect drainage characteristics result in prolonged waterlogging periods during the rainy seasons. In this area, it is common to find patches without crop growth, as observed at site 1. The soil’s physico-chemical properties of each site are consistent with the observations made through the years in this region. Although conservationist management practices were employed, the cultivated crops in these soil types exhibit reduced yields compared to those grown in soils without elevated levels of exchangeable sodium. The decrease in yield can be attributed to the uneven growth of crops, a consequence of the qualities of degraded soil. This is often linked to one or more adverse factors, such as elevated soil pH, electrical conductivity (EC), and salinity (Rengasamy et al., 2022). On the other hand, considering that the community structure of AMFs depends on chemical and physical properties of soil, it is interesting to explore the indigenous taxa adapted to adverse conditions as a potential strategy to increase crop production in this region. Thus, the aim of this study was to characterize native arbuscular mycorrhizal fungal communities in these field sites. To the best of our knowledge, this is the first report of AMF communities description based on sequencing of rRNA amplicons using the ONT. Many previous reports of AMF communities were based on short read sequencing, mainly Illumina sequencing (Higo et al., 2020; Stefani et al., 2020; Loit et al., 2023). The amplicon used in these studies targeted the ITS region of AMF, with sizes ranging between 300 bp (AMV4.5NF and AMDGR) and 550 bp (NS31 and AML2). Few studies used long read sequencing, mainly Pacific Biosciences sequencing (Schlaeppi et al., 2016; Kozjek et al., 2021; Kolaříková et al., 2021), allowing to obtain longer reads that spanned the region containing the 18S rRNA gene, ITS and part of the 28S rRNA gene. Having longer sequences, allows for a better taxonomic resolution than short reads (Krehenwinkel et al., 2019). One drawback from ONT is that, in flowcells 9.4.1, the error rate is significantly higher than that of Illumina and PacBio, which may make the bioinformatic analysis of amplicons more difficult. However, using a clustering and polishing approach, errors can be corrected, and clusters can be obtained to perform ecological analysis. There are few reports of amplicon sequencing that used this approach (Beale et al., 2022; Maes et al., 2023), but with new bioinformatic tools and improved flow cell chemistry this trend will continue to grow. In this work, we used a clustering approach using abPOA, which works faster than other algorithms like sPOA (Vaser et al., 2017), and improved cluster quality with medaka to minimize errors. Moreover, to improve results, we discarded all clusters with unexpected lengths and those that lacked one or both primers. Another drawback of using long reads is that databases like MaarjAM (Öpik et al., 2010) are based mainly on small sequences. Therefore, in many cases only a small fragment of the resulting cluster is used for taxonomic profiling. Nevertheless, long clusters are better when comparing with longer sequences, like the Kruger fragment (∼1600 bp), and for phylogenetic analysis. More studies based on long reads would improve taxonomic profiling of AMF.

Our research not only involved the assessment of innovative tools for characterizing mycorrhizal fungal communities but also provided valuable insights into their potential adaptation to soil degradation given by alkalinity and salinity conditions. Previous studies, using Illumina sequencing, observed variations in AMF communities (OTUs, based on a sequence similarity threshold of 97%) along a gradient of soil salinity and arsenic contamination, but also associated to: soil pH, moisture, organic matter content and plant available soil phosphorus (Parvin et al., 2019). Moreover, studies in a salinized south coastal plain in China showed that abiotic factors (soil salinity and pH) and host plant traits (salt tolerance, lifestyle or origin) determine the composition and distribution of root-colonizing AMF taxa in the salinized ecosystem (Guo & Gong 2014). In this study, sequencing results did not reveal significant differences neither in diversity nor abundance of the AMF community among the analyzed soil samples. Despite the influence of soil parameters on the variation in spore densities across diverse production areas, it is noteworthy that elevated levels of diversity and abundance do not consistently align with mycorrhization frequencies between different production areas (Houngnandan et al., 2022).

Our results showed that the *Glomeraceae* family was present in a greater proportion in all the evaluated sites, demonstrating its great adaptation to salinity and alkaline pH. The *Acaulosporaceae* family emerged as the second most prominent family, with *Gigasporaceae* and *Paraglomeraceae* following to a lesser degree. *Glomeraceae* represented between 43 and 95% of the AMF community in the studied field sites. Within the Glomeraceae family the most representative genus in all the soils samples were: *Glomus, Rhizophagus, Septoglomus* and *Funneliformis*.

The dominance of the *Glomeraceae* family in stressful habitats of Argentina has already been reported in Becerra et al., 2019. Likewise, Stürmer et al., (2018) found that Glomereaceae is one of the two families that dominate maritime sand dunes worldwide, and its relative species richness positively correlates with soil pH. The proportion of species in Glomeraceae exceeded 40% in soil with pH > 6.5 (Stürmer et al., 2018). The dominance of *Glomeraceae* has been observed in soils as well as in roots subjected to different environmental conditions (Akaji et al., 2022; Kaidzu et al., 2020; Faggioli et al., 2019; Borriello et al., 2012; Öpik et al., 2010).

In alkaline soils, the number of sequences of the genus *Destiscutata* (from the *Gigasporaceae* family) and unclassified members of the *Glomeromycetes* class increases. There is also a slight increase in members of the *Acaulospora* genus (*Acaulosporaceae*). Notably, alkaline saline sodic soils exhibited the highest amount of *Acaulospora* and the loss of members of the genus *Septoglomus* was observed.

*Acaulosporaceae* was reported as the dominant family in several harsh conditions (i.e., contaminated soils) and have demonstrated advantages for host plants protection in polluted sites (Morton 1986; Barbosa et al., 2017; Garcés‐Ruiz et al., 2019; Kaur & Garg 2022; Wang et al., 2007). In contrast to prior studies, which linked *Acaulospora* with acid soils and low soil EC (Morton 1986; Veresoglou et al., 2013; Vieira et al., 2020; Akaji et al., 2022), our research reveals a broader adaptability. Similarly, Garcés‐Ruiz et al., (2019) detected that *Acaulospora* thrives not only in alkaline soils with a pH 8 but also in environments with lower pH levels ranging from 5 to 6, showcasing its remarkable versatility. On the other hand, an inhibition of *Acaulospora* spore germination has been observed in saline environments (Juniper & Abbott 2006). Nevertheless, *Acaulosporaceae* species establish symbiotic relationships with mangrove trees, potentially enhancing the nutritional status of these plants in saline and nutrient-deficient settings (Akaji et al., 2022). Hence, although *Glomus* and *Acaulospora* seem to be the predominant native AMFs in ASS soils, further studies should investigate if the remodeling of the community structure alleviates the limitation in plant growth under these adverse conditions safeguarding crop production and environmental sustainability.

## CONCLUSION

In this study, we examined the arbuscular mycorrhizal fungal (AMF) community across three distinct sites with varying soil properties. Our research is among the first to describe AMF communities by sequencing a large fragment of *Glomeromycota* rDNA. Despite the challenges posed by moderate error rates in ONT flowcells (R9.4.1. chemistry), our clustering and polishing approach demonstrated effective error correction and cluster generation for ecological analysis. Our results indicate no significant differences in AMF community diversity and abundance among soil samples. Nevertheless, our findings emphasize the potential of *Glomus* and *Acaulospora* in ASS soils, with further exploration needed to understand how these AMFs contribute to plant growth under adverse conditions. The study not only contributes to the characterization of native AMF communities but also highlights the importance of innovative sequencing tools in advancing our understanding of these vital symbiotic relationships.

## ACKNOWLEDGMENT

This research was funded by the National Institute of Agricultural Technology of Argentina (2023-PE-L04-I073).

## CONTRIBUTIONS

CMB and GML conducted the field sampling.MN, FFD and GML performed laboratory work. IJM AAF and FFD assisted with bioinformatics data analysis. MN interpreted the data. MN, IJM and GML wrote the first draft of the manuscript. All authors contributed to the final version of the manuscript.

## DECLARATION OF COMPETING INTEREST

The authors declare that they have no known competing financial interests or personal relationships that could have appeared to influence the work reported in this paper.

## FIGURES

**Supplementary Table 1:**
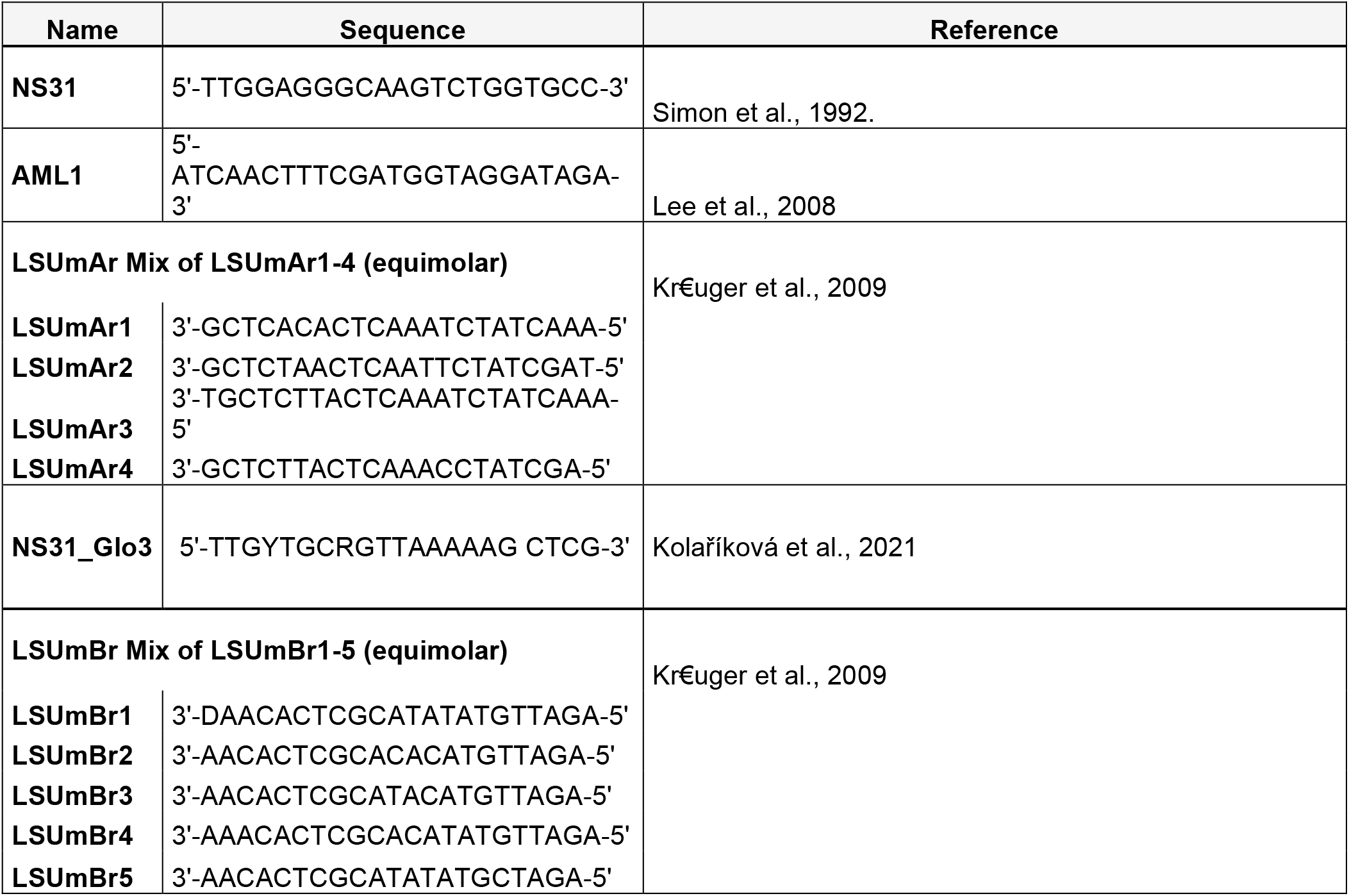
List of AMF-specific primers.

## Notes

### Competing Interest Statement

The authors have declared no competing interest.

### Summary of Updates

Previous Title: "Analysis arbuscular mycorrhizal fungi (AMF) community composition associated with alkaline saline sodic soils" was changed to "Analysis of arbuscular mycorrhizal fungi (AMF) community composition associated with alkaline saline sodic soils"

